# Can we test the influence of prosociality on high frequency heart rate variability? A double-blind sham-controlled approach

**DOI:** 10.1101/078006

**Authors:** Brice Beffara, Martial Mermillod, Nicolas Vermeulen

**Author notes:** Correspondence concerning this article should be addressed to Brice Beffara, Office E250, Institut de Recherches en Sciences Psychologiques, IPSY - Place du Cardinal Mercier, 10 bte L3.05.01 B-1348 Louvain-la-Neuve, Belgium.

## Abstract

The polyvagal theory (Porges, 2007) proposes that physiological flexibility dependent on heart-brain interactions is associated with prosociality. So far, whether prosociality has a causal effect on physiological flexibility is unknown. Previous studies present mitigated results on this matter. In a randomized double-blind protocol, we used a generation of social closeness procedure against a standardized control condition in order to manipulate social affiliation as a prosocial interaction factor. High frequency heart rate variability (HF-HRV, indexing physiological flexibility), electromyographical activity of the corrugator supercilii (sensitive to the valence of the interaction) and self-reported measure of social closeness were monitored before, during, and after experimental manipulation. Cooperation was measured after the experimental manipulation as an index of behavioral prosociality. Data reveal no evidence toward and effect of the experimental manipulation on these measures. We discuss methodological aspects related to the experimental constraints observed in social psychophysiology. Implications for the experimental test of the polyvagal theory are approached within alternative theoretical frameworks.

## Introduction

Prosocial interactions are associated with positive health and well-being states (S. L. Brown & Brown, 2015). More specifically, affiliate behaviors play an important part in coping with stressful events (Raposa, Laws, & Ansell, 2015). According to the polyvagal theory (Porges, 2007), heart-brain interactions are a central mechanism in the interplay between stress and prosocial behaviors. Particularly, efficient activity of the myelinated vagus nerve connecting the heart and the brain is proposed to foster capacities fully required when the organism has to adapt to external demands and internal needs such as during social interactions (Taborsky & Oliveira, 2012). This adaptability is referred as physiological or autonomic flexibility (Brosschot & Thayer, 1998; B. H. Friedman & Thayer, 1998; Thayer & Lane, 2000) in reference to the ability of the organism to show dynamic variations in response to the continuous variations of the environment. In the social domain, it has indeed been shown that physiological flexibility was associated with prosociality (Beffara, Bret, Vermeulen, & Mermillod, 2016; J. G. Miller, Kahle, & Hastings, 2015).

Even if important limitations have been suggested toward the polyvagal theory (Grossman & Taylor, 2007; Taylor et al., 2014), it remains that more and more evidence corroborate the predicted link between the myelinated vagal functioning and affiliative social tendencies (Bornemann, Kok, Böckler, & Singer, 2016; Kogan et al., 2014; Muhtadie, Koslov, Akinola, & Mendes, 2015). What remains unclear, however, is the direction of this association. Indeed, better heart-brain interactions could lead to improved social skills, or conversely, or even a third factor could link these two variables. Kok & Fredrickson (2010) proposes that the association is actually bidirectional and that myelinated vagal activity and social experiences reciprocally influence each other in a dynamic loop. This proposition is highly relevant since the development of social skills likely depends on progressive learning processes interacting with the evolution of cardiovascular regulation (Brosschot, Verkuil, & Thayer, 2016b, 2016a). However, the method of Kok & Fredrickson (2010) has been recently criticized, both on statistical and physiological aspects (Heathers, Brown, Coyne, & Friedman, 2015) but these criticisms have been adequately answered (B. E. Kok & Fredrickson, 2015). The proposition remains important as it allows new hypotheses formulations in the testing of the causal link between the quality of heart-brain interactions and social functioning. However, to our knowledge, paradigms allowing to test the causal direction in an experimental way have not been reported yet.

A recent meta-analysis (Shahrestani, Stewart, Quintana, Hickie, & Guastella, 2015) on this matter concludes that positive social interactions do not increase myelinated vagal functioning but negative interactions decrease it. One given explanation is that positive interactions could be beneficial after stressful events but not in a context already “favorable”, which is in line with previous propositions (Raposa et al., 2015). The meta-analysis was performed on 14 studies including 17 tasks, among which 10 deal with negative social interactions (stressful), 3 with neutral interactions, and only 4 with positive interactions. Looking more closely at the 4 positive social tasks, we can observe that the manipulation of the valence of the social task in the positive direction leads hardly to conclude that a modulation of the affiliative functioning of the dyad actually happens.

Among the 4 studies dealing with positive interactions, the first reported in the meta-analysis has been carried-out by Butler, Wilhelm, & Gross (2006). Their experimental manipulation focused on an emotion regulation instructional set concerning a negative film. This instructional set was delivered for only one of the two members of the dyad. As a consequence, although the valence of the task was manipulated, affiliation was not technically central in their experimental design. The experimental manipulation of D’Antono, Moskowitz, Miners, & Archambault (2005) was closer to a form of social closeness generation using agreeable versus quarrelsome role-play in dyads. However, as the prosocial nature and the effect (S. L. Brown et al., 2009) of agreeable role-play was not assessed, it is hard to determine whether myelinated vagal functioning was not influenced by provoked prosociality, or if prosociality was not actually successfully induced. On the contrary, Kathi J Kemper & Shaltout (2011) seem to find an effect of prosociality on autonomic flexibility but the protocol includes non-verbal communication techniques which necessarily add confounding factors to the manipulation of more natural affiliative social behaviors. What is more, sample size was very small (n=5) and the study was carried out in a healthy volunteer-clinician dyad, which did not allow applying the double blind during the experimental manipulation. Finally, because the sample size reported in Willemen, Goossens, Koot, & Schuengel (2008) is larger, the increased autonomic flexibility observed after a positive social interactions is much more reliable. Nonetheless, the main methodological features of the experimental design do not allow to fully conclude to an effect specific to the prosocial interaction. Indeed, the study involves adolescent-parents interactions after a stressful event. The aim of the study was to determine the effect of the parent visit on stress recovery of the adolescent. As no control group was set up, there is still a possibility that the mere presence of another individual would have resulted in physiological modifications.

Collectively, this set of 4 studies gives important clues about the potential effect (or absence of effect) of prosocial interactions on autonomic flexibility. Despite all these efforts, we believed that the issue could be addressed by the mean of a complementary methodological design (S. L. Brown et al., 2009).

We used an experimental design based on the work of A. Aron, Melinat, Aron, Vallone, & Bator (1997). This protocol enables to generate social closeness by the mean of guided discussions in dyads through short sentences such as questions or instructional sets. The content of these phrases promote the exchange of information between the two persons in the dyad. This exchange of information is expected to provoke reciprocal self-disclosure and engage the two persons in a prosocial interaction by sharing autobiographical elements about themselves. This has been shown to increase subsequent altruism (S. L. Brown et al., 2009). As a consequence, this design is particularly appropriate to test the polyvagal theory in the “social to physiology” direction. We also set up a combination of apparatus permitting to blind the experimenter to the condition of the participants (social closeness or a control condition also developed as a neutral “small talk” condition by A. Aron et al. (1997)).

As compared to several studies included in Shahrestani et al. (2015), we operationalized autonomic flexibility as the high frequency component of heart rate variability (HF-HRV, the variation in the cardiac beat to beat intervals, Heathers (2014)). HF-HRV is a reliable and noninvasive measurement of the dynamics of short-term heart-brain interactions (Thayer, Åhs, Fredrikson, Sollers, & Wager, 2012). We also measured the electromyographical activity of the corrugator supercilii muscle (involved in frowning and emotional facial expressions of anger) as a secondary measure of autonomic activity. The corrugator supercilli activity is increased by negative and decreased by positive valence (J. T. Larsen, Norris, & Cacioppo, 2003), which should be generated by our social closeness condition. Moreover, the corrugator supercilii is sensitive to threat (Costa, Bradley, & Lang, 2015), which should be diminished by our social closeness condition. Both HF-HRV and corrugator supercilii activity should then be able to measure the effect of social closeness manipulations following the proposition of the polyvagal theory (Porges, 2007).

Finally, our design also includes a task of cooperation after the social closeness generation (or control) condition in order to evaluate whether manipulating prosociality as affiliation and interpersonal positive contact can transfer to a behavioral measure of prosociality. A dot detection task in dyads was used, where the participant must try to press a key after a signal, simultaneously with the other participant (X. Cheng, Li, & Hu, 2015; Cui, Bryant, & Reiss, 2012). Cooperation is expected to increase response times in order to synchronize with the other participant while reducing the time between the response times of the two participants.

We hypothesized that, compared to a control condition, social closeness generation should increase HF-HRV, decrease the activity of the corrugator supercilii, and increase cooperation. We also predict that this effect should be mainly observed in low baselines participants for HF-HRV and high baseline participants for EMG activity. Therefore, social closeness would benefit more to participants with lower autonomic flexibility and higher default stress response (Brosschot et al., 2016b, 2016a). Indeed, this hypothesis is based on a “deficit remediation” model according to which participants with deficits in a specific parameter will receive more benefits from the manipulation of this parameter (I. W. Miller et al., 2005, 2008). As a consequence lower/higher pre-manipulation HF-HRV/corrugator activation should predict higher level of progression after experimental manipulation. For instance, Davies, Niles, Pittig, Arch, & Craske (2015) showed that cognitive behavioral therapy efficiency was higher for lower pre-manipulation HF-HRV participants, a mechanism we hypothesize to apply also to our manipulation.

## Method

### Sample

Initial sample was composed of 104 healthy human adults. Participants were recruited via advertisements (spread on facebook groups related to Louvain-la-Neuve, Belgium, where the experiment took place). Participants were from the general population and were French speaking. They provided written informed consent before the participation. The study was reviewed and approved by the ethics commission of the Psychological Sciences Institute of the Catholic University of Louvain, Belgium (reference number 15-37). To be eligible, participants had to be aged between 18 and 60 years, with a normal or normal-to-corrected vision, explicitly reported an absence of psychiatric, neurological, hormonal, or cardiovascular disease, and with no medical treatment (with the exception of contraception). Smoking, energizing drinks (e.g. coffee, tea, etc…) and psychotropic substances (e.g. alcohol, cannabis, etc…) were prohibited to each participant the day of the experiment. They had also to avoid eating or drinking (water was allowed) the 2 hours preceding the experiment in order to limit the influence of digestion on autonomic functioning (Short term HRV measurement can be biased by the digestion of food since viscera are innervated by the autonomic nervous system, (Heathers, 2014; Iorfino, Alvares, Guastella, & Quintana, 2016; Quintana & Heathers, 2014)) but they had to eat in the morning (more than 2 hours before the experiment) in order to avoid fasting states. The participants received experimental 15 euros at the beginning and 20 euros at the end of the period the recruitment in order to complete our sample.

### Sample size

We planned one hundred and sixty participants to take part in the study in order to work with a similar sample size as compared to S. L. Brown et al. (2009). Their sample size was adequate to observe an effect of an experimental generation of social closeness on progesterone compared to a neutral control task, with an effect size of *R*^2^~.63. Unfortunately, even with an increase of the compensation, we could not reach this sample size.

### Procedure

After completing the inclusion survey online, participants suitable for participation were automatically redirected toward another survey in order to give their available dates for an appointment (others were thanked and informed that they did not fit with the criterion). A homemade R-script was built in order to randomly select dyads among all participants available at each slot. The appointment date and time was determined and communicated to the participants roughly 72h before the actual slot. The experiment took place in a quiet and dimmed room. All participants were tested between 0900 h and 1300 h. Participants were asked to go empty their bladder before starting the experiment. After a global description of the experiment, they were taught how to install the Bioharness^®^ heart rate monitor. They were left in autonomy in an isolated room for the installation of the heart rate monitor. Then, they seated in a chair, the experimenter checked the signal and the installation of EMG electrodes began. The three electrodes (two on the corrugator supercilii and on the top of the forehead) were attached and the signal was checked. Classical piano music (Ballade No.4 in F minor, Op. 52 by Frederic Chopin, interpreted by Franck Levy https://musopen.org/fr/music/769/frederic-chopin/ballade-no-4-op-52/) was played during the installation. We added this feature in order to compensate for the potential stressful effects of electrodes’ installation. The experiment started when the quality of the signals were correct.

First participants had to perform facial actions in order to get a baseline of the volitional contraction of the corrugator supercilli. They had a succession of 2*10=20 instructional sets randomly displayed on their computer screen: “Frown then relax”, “Swallow”, “Wrinkle your eyes then relax”, “Clench the jaws then relax”, “Close then open your eyes”, “Close then open your mouth”, “Raise the corner of your lips then relax”, “Raise your eyebrows then relax”, “Wrinkle your nose then relax”, “Lower the corner of your lips then relax”. The “Frown then relax” instructional set was used to compute maximum possible signal level and other instructions had a distraction role in order to avoid too much focusing on the frowning action during the experiment. Instructions were displayed for 3 seconds on the screen and followed by a 3 second new instruction to relax the face.

In a second step, participants were asked to answer some questions about their relationship with the their partner (i.e. the other participant of the dyad, (A. Aron & Fraley, 1999; A. Aron et al., 1997)). First, they were asked if they knew their partner on a Likert scale from 1 = “Not at all” to 7 = “Perfectly well”. Second, they were asked how close they felt to their partner on a Likert scale from 1 = “Not at all”, from 7 = “Enormously”.

During the 5 following minutes, participants watched short neutral samples of films selected and evaluated by Hewig et al. (2005) (“Hannah and her Sisters” and “All the President’s Men”) and Schaefer, Nils, Sanchez, & Philippot (2010) (“Blue [1]”, “Blue [2]”, “Blue [3]” and “The lover”). Videos were displayed without audio. These first 5 minutes aimed to allow participants to shift in a resting state. ECG data for HRV baseline computation was recorded for the 5 following minutes while participants listened to the first 5 minutes of a neutral audio documentary designed for laboratory studies (Bertels, Deliens, Peigneux, & Destrebecqz, 2014). Neutral videos and audio documentary were used in order to standardize ECG recordings (Piferi, Kline, Younger, & Lawler, 2000).

After resting-state recording, participants were put in a discussion situation for a minimum of 10 minutes. The protocol is detailed below. Another resting-state recording was performed after the discussion for 5 minutes while participants listened to the last 5 minutes of the neutral audio documentary (Bertels et al., 2014). Then participants performed the cooperation task detailed below. The last step of the experiment in laboratory was again a question about how close participants felt to their partner on a Likert scale from 1 = “Not at all”, from 7 = “Enormously”. EMG electrodes were detached (classical music was played), the ECG belt uninstalled, and a debriefing was proposed before compensating the participants. Control survey was completed at home, online, on Qualtrics, thanks to an identifier given to the participants. ECG data was recorded during spontaneous breathing (Denver, Reed, & Porges, 2007; Kobayashi, 2009; Kowalewski & Urban, 2004;P. D. Larsen, Tzeng, Sin, & Galletly, 2010; Muhtadie et al., 2015; Pinna et al., 2007). The experimenter was available at any time during the experiment but stayed in another room.

### Generation of social closeness

Dyads were randomly and automatically assigned to the closeness generation condition or control condition. Participants were blind to their condition, such as the experimenter. Both conditions were guided discussions where participants took turn asking a question to their partner –and the partner had to answer the question– or following a small instruction in order to engage in conversation. The phrases used to guide the conversation were taken from A. Aron et al. (1997), translated in French by external translators (3 pairs of translation and back-translation) and reviewed, adapted, and selected by us. The minimum time for discussion was 10 minutes, but could last a bit more depending on the duration of the last item (phrase). Thirty-six items were available for the discussion which was largely enough to fill 10 minutes, even if participants move quickly from one item to another. Items were displayed sequentially and the participants chose to move to the following item. Items were used for each participant (participant 1 asks participant 2 and vice versa). In the social closeness condition, phrases are designed to foster, as described by A. Aron et al. (1997) “sustained, escalating, reciprocal, personalistic self-disclosure” (A. Aron et al., 1997, p. 364) such as “Given the choice of anyone in the world, whom would you want as a dinner guest?”, “When did you last sing to yourself? To someone else?”, or “Is there something that you’ve dreamed of doing for a long time? Why haven’t you done it?”. In the control condition phrases are more neutral and less likely to engage this kind of process, such as “When was the last time you walked for more than an hour? Describe where you went and what you saw.”, “If you could invent a new flavor of ice cream, what would it be?”, “Where are you from? Name all of the places you’ve lived.”.

During all the experiment, the participants were seated opposite to each other but separated by a panel such that they could not see each other. Each participant had his/her own screen, and the two screens were connected to the same computer. A webcam was attached to each screen and was automatically activated at the beginning of the discussion phase and shut down at the end of the same phase. The video from the webcam was displayed in real time on the screen of the other participant. As a result, participants could see each other only during the discussion phase by the mean of the webcam. We programmed, using the OpenCV library for Python, the automatic management of the web-cams and their coordination. The script was integrated in the Psychopy 1.8 script in order to match the progress of the experiment. Web-cams were calibrated at the beginning of each testing in order to center the image on the face of the participant.

\#### Cooperation We used the cooperation task described in Cui et al. (2012) and X. Cheng et al. (2015) in order to evaluate behavioral prosociality after experimental manipulation. This task includes a cooperation task, competition task and a neutral task. The competition and neutral tasks serve as control tasks. Each trial begins with a hollow gray circle at the center of the screen of each participant that stays visible for a random interval between 0.6 and 1.5 s. Subsequently, a green cue signals the participants to press a response key (left arrow for one and right arrow for the other. A green sticker was attached to the keys in order to make them salient on the keyboard). During a training phase of 5 trials, participants had to be relatively constant in their response times. Their response time was displayed as a feedback after the first trial, and then the feedback was “+ 1 point” if they succeeded in being constant or “−1 point” if they failed for the following trial. A response time was determined as constant compared to the previous trial if the difference between the two trials was inferior to a threshold T = (RT1+RT2)/10 (adapted from Cui et al. (2012)). The aim of the participant was to get a maximum number of points for each phase. The score was reset to 0 at the end of each phase.

The first phase of interest was the phase of cooperation composed of 20 trials. Participants had to coordinate (without talking to each other) in order to press the key simultaneously. A trial was successful if the difference of response times between the two players was inferior to T = (RT1+RT2)/10. Again, a feedback “+ 1 point” was displayed if they succeeded in cooperating or “−1 point” if they failed.

The second phase was the competition phase where the aim was to press the key faster than the other player. The fastest player won 1 point and the slowest lost one. The third and last phase was played alone with the same instructions as the training phase detailed above.

The points were uniquely related to the task and had no other consequences on the experiment. Each negative feedback (“−1 point”) was associated with the display of the difference between response times. In the “competition” phase, the difference between response times was always displayed alongside the points. The task was coded in Python.

## Physiological measurement

### Electrocardiography

The electrocardiogram (ECG) data was recorded with a Zephyr Bioharness^TM^ 3.0 (Zephyr, 2014). The Bioharness^TM^ is a class II medical device presenting a very good precision of measurement for ECG recording in low physical activity conditions (Johnstone, Ford, Hughes, Watson, & Garrett, 2012a, 2012b; Johnstone et al., 2012). It has been used for ECG measurements in both healthy and clinical populations, presenting a very high-to-perfect correlation with classical hospital or laboratory devices (Brooks et al., 2013; Yoon, Shah, Arnoudse, & De La Garza, 2014). The BioharnessTM both provides comfort for the participant and allows reliable HRV extraction for the researcher (Lumma, Kok, & Singer, 2015). The chest strap’s sensor measures electrical activity corresponding to the classical V4 lead measurement (5th intercostal space at the midclavicular line) through conductive Lycra fabric. A single-ended ECG circuit detects QRS complexes and incorporates electrostatic discharge protection, both active and passive filtering and an analog-to-digital converter. Interbeat intervals are derived by Proprietary digital filtering and signal processed with a microcontroller circuit. The ECG sensor sampling frequency is 250 Hz and the resolution 0.13405 mV, ranging from 0 to 0.05 V (Villarejo, Zapirain, & Zorrilla, 2013). After a slight moistening of the 2 ECG sensors, the chest-strap was positioned directly on the skin, at the level of the inframammary fold, under the lower border of the pectoralis major muscle. The recording module communicated with an Android^®^ OS smartphone by Bluetooth^®^. The application used to acquire the signal emitted by the Bioharness^TM^ was developed, tested, and validated by Cânovas, Domingues, & Sanches (2011). The Android^®^ OS device used to record the signal was an LG-P990 smartphone (Android^®^ version 4.1.2.).

### Electromyography

EMG was measured with three 4 mm Ag/AgCl electrodes: two electrodes were attached at the level of left brow (central part, just above the brow) and one ground sensor was placed upon the participant’s top left forehead (Fridlund & Cacioppo, 1986). Sampling frequency was set at 2000 Hz.

### Control for confounding factors

To control for confounding variables likely to be linked to HRV, participants completed questionnaires detailing life habits, demographic data and emotional traits (Quintana, Guastella, Outhred, Hickie, & Kemp, 2012). Physical activity was assessed with the International Physical Activity Questionnaire (IPAQ,Craig et al. (2003)), composed of 9 items that calculate an index reflecting the energy cost of physical activities (Metabolic Equivalent Task score, MET). The IPAQ has been validated in French (Briancon et al., 2010; Hagströmer, Oja, & Sjöström, 2006) and widely used in French surveys (Salanave et al., 2012). Participants also completed the Depression Anxiety and Stress scales (DASS-21;(P. F. Lovibond & Lovibond, 1995)). The DASS-21 is a 21-item questionnaire, validated in French (Ramasawmy & Gilles, 2012), and composed of three subscales evaluating depression, anxiety and stress traits. We also recorded the size, weight, age and sex of the participants and their daily cigarette consumption. Participants answered final surveys at home via an online survey built with Qualtrics in order to reduce the time spent in the laboratory and to allow all the dyads to be tested between 0900 h and 1300 h.

## Physiological signal processing

### Electrocardiography

R-R interval data was extracted from the Android^®^ device and imported into RHRV for Ubuntu (Rodríguez-Liñares et al., 2011). Signal was visually inspected for artifact (Prinsloo et al., 2011; Quintana et al., 2012; Wells, Outhred, Heathers, Quintana, & Kemp, 2012). Ectopic beats were discarded (Kathi J. Kemper, Hamilton, & Atkinson, 2007) for participants presenting a corrupted RR interval series (Beats per minute (bpms) shorter/longer than 25/180 and/or bigger/smaller than 13% compared to the 50 last bpms). RR series were interpolated by piecewise cubic spline to obtain equal sampling intervals and regular spectrum estimations. A sampling rate of 4 Hz was used. We then extracted the frequency component of HRV from RR interval data. The LF (0.04-0.15 Hz) and HF (0.15-0.4 Hz) components were extracted using an east asymmetric Daubechies wavelets with a length of 8 samples. Maximum error allowed was set as 0.01 (García, Otero, Vila, & Márquez, 2013).

### Electromyography

Signals were re-sampled at 1000 Hz, amplified, filtered through a 30–250 Hz band pass and 60 Hz notch, digitized, re-filtered, rectified, and then integrated over 20 ms (Bershad, Seiden, & Wit, 2016) online using EMG100C amplifiers, an MP150 data Acquisition System, and Acqknowledge software from Biopac Systems (Goleta, CA, USA). Maximum acceptable amplitude of the signal was computed as the maximum signal value during the volitional contraction of the corrugator supercilii (frowning). All values superior to this threshold were reduced as the threshold value. EMG files presented a median [interquartile range] of 0.00 [0− 0.01] % of ectopic EMG values with a maximum value of 2.13 %.

## Data analysis

### Data preparation

Excluding technical problem recordingsˆ[Numerous recording problems happened notably due to frequent freezing of Psychopy when managing the synchronization of the two web-cams. Moreover, a lot of participants lost their identifier and did not complete the control questionnaires, we therefore excluded these data for final analysis for all participants. Here we present the results concerning a restricted part of our sample where all data (excepted demographic and self-reported at home) are available, a total of 72 participants were available for data analyses (Figure 1). 4 participants were excluded from the sample before data analysis because of a noisy ECG signal. Analysis on physiological signals were performed on data averaged on 5 minutes for each experimental step (t1= resting baseline, t2 = first part of the discussion, t3 = second part of the discussion, t4 = resting after discussion). As a result, 4 measurement points of 5 minutes were available for 2 physiological measurements (HF-HRV and EMG_*corrugatorsupercilii*_). All participants included in final analysis presented a median [interquartile range] of 0.03 [0 – 0.16] % of ectopic beats values with a maximum value of 8.17 %. We computed the HF power of HF-HRV baseline and then calculated as the natural logarithm of the HF power in order to correct the right-skewed distribution (Kogan et al., 2014; Pinna et al., 2007). In order to correct for the positive skew of EMG data, data were square-root transformed (J. T. Larsen et al., 2003) but normality could not be reached for each time step. Behavioral data in the cooperation task (measured after t4) were analyzed as raw reaction times and difference of reaction times between participants of the dyad as a function of the experimental group.

**Figure 1.**
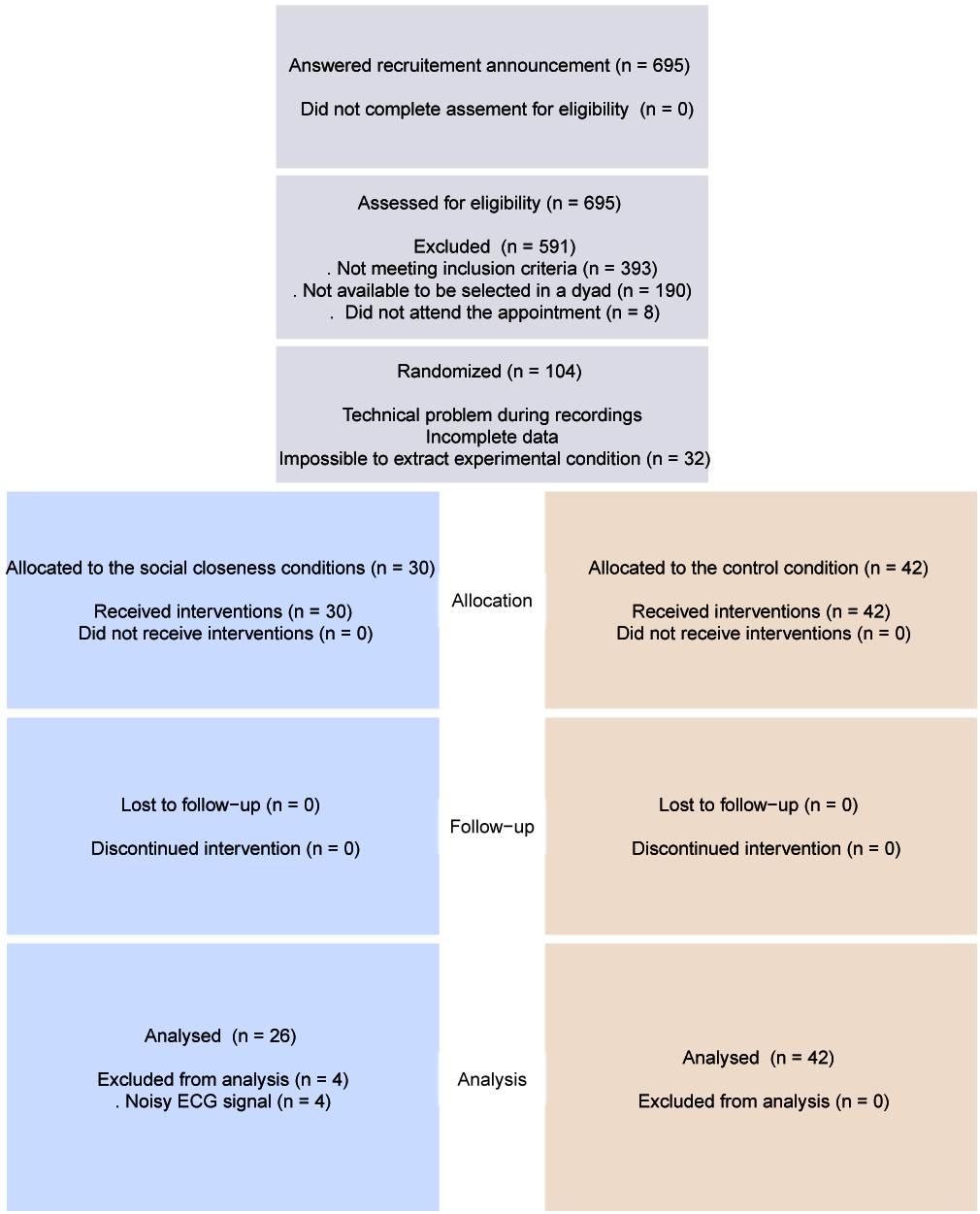
Flow chart of participants

### Model comparison

Statistical analysis were conducted using RStudio^®^, version 1.0.8 for Linux (R Core Development Team, 2015) and are reported with the knitr (Xie, 2013), papaja (Aust & Barth, 2015) and rmarkdown (Allaire et al., 2016) packages. The aim of data analysis was to detect whether experimental social closeness generation (compared to the control condition) influenced physiological and behavioral variables across time and whether this effect was dependent on the baseline level of participants. We compared alternative models (for alternative hypothesis = “H1”) with experimental group (called “G”: Closeness vs. Control) and models with experimental group and resting baseline (called “R” = t1) as an independent variable to a model containing only the intercept (Null hypothesis = “H0” model) with signals at each time step and channel as dependent variables. Behavioral data measured during the cooperation task at the end of the experiment (after t4) were analyzed as a function of the experimental group and task type (competition session, alone situation session, and cooperation session, respectively coded as −1, 0, 1).

We analyzed our data by the fit linear mixed-effects models function (lmer), computed using the package “lme4” [Bates et al. (2014);Bates2015] for behavioral data and linear models function (lm) for physiological data (Chambers, 1992). All residuals of the models were not normally distributed but data transformation did not allow getting normal distribution. Model selection was completed using AICc (corrected Akaike information criterion) and Evidence Ratios -*ER*_*i*_-(K. P. Burnham & Anderson, 2004; Kenneth P. Burnham, Anderson, & Huyvaert, 2011; Hegyi & Garamszegi, 2011; Symonds & Moussalli, 2011). In this perspective, all the hypotheses are considered equally, meaning that the status of H0 (absence of effect) is the same as compared to H1 (effect), all models can be compared together. AIC provides a relative measure of goodness-of-fit but also of parsimony by sanctioning models for their numbers of parameters. AICc is more severe on this last point than AIC 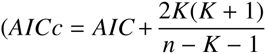 where *K* is the number of parameters and *n* the sample size.). We computed the difference between best (lower value of AICc) and other AICcs with Δ_*AICc*_ = *AICc*_*i*_ − *AICc*_*min*_ thanks to the piecewiseSEM package (Lefcheck, 2016). The weight of a model is then expressed as 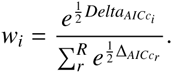. From there, we can compute the Evidence Ratio between the alternative model and the intercept model: 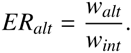. For each physiological and behavioral measurement, we were able to compare the effect of the group and the interaction effect between the intervention group (“G”) and the baseline (“R” = t1 for physiological measurement, only for physiological data) of the participants with the “H0 model” including only the intercept. Baselines were set as continuous factors (for physiological data only) and experimental groups (group: Closeness vs. Control) were coded as +0.5 and −0.5 respectively. If the alternative model (for H1) is more parsimonious than the intercept model (for H0) then substantial (3.2 < *ER* < 10), strong (10 < *ER* < 100) or even decisive (100 < *ER*) evidence should be observed (Kass & Raftery, 1995; Snipes & Taylor, 2014). On the contrary, substantial (1/3.2 < *ER* < 1/10), strong (1/10 < *ER* < 1/100) or even decisive (1/100 < *ER*) evidence toward the intercept model would allow concluding that the intercept model is more parsimonious. An 1/3.2 < *ER* < 3.2 do not allow to draw conclusions and indicates that the data does not provide significant evidence toward one model or the other (Figure 2).

**Figure 2.**
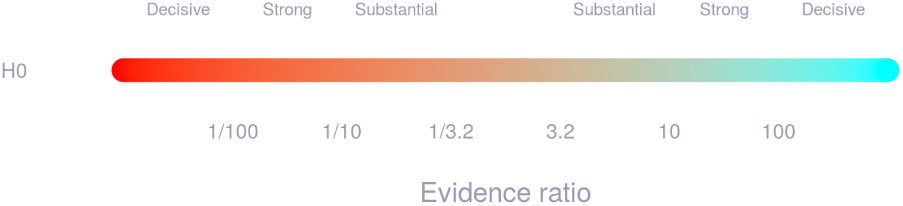
Interpretation of evidence ratios relatively to H0 and H1.

## Results

We first analyzed self-reported measures obtained on Likert scales (7 points), measuring how much participants knew the other participant of the dyad before the experiment, and how much they felt close to him/her before and after the experiment. Participants knew each other similarly and also felt equally close before the experiment in each group (Table 1). Indeed, evidence ratios (<3.2) do not permit to conclude to a difference between groups. The scores between groups were neither different after the experiment, hwever there was strong evidence toward an increase of social closeness in both groups (ER > 100 toward the intercept model compared to 0). Looking at physiological data (Table 2), we did not observe substantial differences between the two conditions (Table 3). Contrary to our hypothesis, there was no evidence toward an interaction between the experimental condition and the baseline. Indeed, all evidence ratios were inferior to 3.2 and, contrary to our hypothesis, do not allow to conclude neither to an effect of social closeness manipulation nor to an interaction with initial physiological levels.

**Table 1.** Effect of experimental group on self-reported measures compared to the intercept model. ER = evidence ratio, SC = Social closeness group, CT = Control group.

|  | ER | SC<sub>mean</sub> | SC<sub>sd</sub> | CT<sub>mean</sub> | CT<sub>sd</sub> |
| --- | --- | --- | --- | --- | --- |
| **Feeling of knowing (before)** | 0.39 | 1.04 | 0.2 | 1.07 | 0.26 |
| **Feeling of closeness (before)** | 1.97 | 1.54 | 0.81 | 1.98 | 1 |
| **Feeling of closeness (after)** | 0.35 | 3.35 | 1.38 | 3.43 | 1.27 |
| **Feeling of closeness (after - before)** | 0.64 | 1.81 | 1.39 | 1.45 | 1.17 |

**Table 2.** Descriptive statistics for HF-HRV (expressed as ms^2^) and EMG data (expressed as µV.s) at each time step of the experiment for each experimental condition.

|  | Measure | t1<sub>mean(sd)</sub> | t2<sub>mean(sd)</sub> | t3<sub>mean(sd)</sub> | t4<sub>mean(sd)</sub> |
| --- | --- | --- | --- | --- | --- |
| **Closeness** | HF-HRV | 5.88 (0.84) | 5.96 (0.87) | 5.92 (0.77) | 6.17 (0.96) |
|  | EMG<sub>corrugator supercilii</sub> | 9.78 (3.78) | 8.98 (2.76) | 8.56 (2.41) | 8.13 (2.43) |
| **Sham** | HF-HRV | 6.15 (0.93) | 6.37 (0.73) | 6.24 (0.83) | 6.36 (0.98) |
|  | EMG<sub>corrugator supercilii</sub> | 10.26 (5.18) | 9.17 (3.82) | 9 (3.62) | 8.43 (3.47) |

**Table 3.** Comparison of alternative models to the intercept model for HF-HRV and EMG data at each time step of the experiment. Reported values are the ER of the alternative model against the intercept model. G = Group, R = resting-state baseline (at t1 measurement).

|  | Factor | t1 | t2 | t3 | t4 |
| --- | --- | --- | --- | --- | --- |
| **HF-HRV** | *G* | 0.70 | 2.79 | 1.19 | 0.46 |
|  | *G \* R* | - | 0.70 | 0.49 | 0.34 |
| **EMG<sub>corrugator supercilii</sub>** | *G* | 0.36 | 0.34 | 0.39 | 0.36 |
|  | *G \* R* | - | 0.36 | 0.44 | 0.38 |

**Table 4.** Descriptive statistics for reaction times and differences of reaction times during cooperation and control tasks expressed in milliseconds.

|  | Measure | Alone<sub>mean(sd)</sub> | Competition<sub>mean(sd)</sub> | Cooperation<sub>mean(sd)</sub> |
| --- | --- | --- | --- | --- |
| **Closeness** | *RT* | 349.93 (114.78) | 293.04 (143.54) | 346.21 (162.04) |
|  | *Diff* | 110.31 (116.69) | 106.35 (193.54) | 91.59 (167.42) |
| **Control** | *RT* | 325.99 (105.32) | 281.4 (182.04) | 356.8 (248.06) |
|  | *Diff* | 82.66 (111.18) | 90.8 (229.05) | 102.32 (268.44) |

**Table 5.** Comparison of alternative models to the intercept model for reaction times and differences of reaction times during cooperation and control tasks. Reported values are the ER of the alternative model against the intercept model. *** indicates strong evidence toward the alternative model (H1).

|  | RT | Difference |
| --- | --- | --- |
| **Experimental group** | 0.51 | 0.81 |
| **Type of task** | >100*** | 0.37 |
| **Group\*Type** | 0.46 | 1 |

The same result appeared concerning behavioral data^1^. Contrary to our hypothesis, there was no evidence toward an effect of group and task type (cooperation, competition, and the single condition) on reaction times, nor on differences of reaction times between participants of a same dyad. Only the task type had a strong effect on reaction times following a linear relationship (competition, alone situation, and cooperation respectively coded as −1, 0, 1, Tables 4) and 5. Again, contrary to our hypothesis, participants in the social closeness condition did not differ from participants in the control condition in terms of reaction times in the dot detection task,either for cooperative, competitive, or alone situation. Experimental manipulation did not modify behavioral response synchronization between participants of a same dyad.

## Discussion

This experiment was designed in order to test the influence of the experimental generation of social closeness on the autonomic nervous system activity. In addition to physiological measures, we evaluated the participants’ level of cooperation. Data collected on self-reported measures, physiological signals (HF-HRV and EMG), and behavioral measures of prosociality (cooperation) did not bring substantial evidence toward an effect of group: social closeness versus control. We expected an interaction with baseline such as low baseline participant on the variable of interest would benefit more from the treatment. Again, this hypothesis was not well supported by the data. As a consequence, data do not allow concluding that a protocol of interpersonal closeness generation can impact physiological flexibility as indexed by HF-HRV and secondary by EMG activity, nor behavioral prosociality as indexed by cooperation. In double-blind sham-controlled conditions, short term positive interpersonal interaction by personal information sharing does not differ from a neutral interaction at physiological, behavioral and self-reported levels. Following the meta-analysis carried out by Shahrestani et al. (2015), we set up a protocol in a randomized double-blind design in order to determine whether or not positive affiliative social interactions could influence the physiological correlates of prosociality (HF-HRV and EMG activity of the corrugator supercilii) and other prosocial skills such as cooperation. A. Aron et al. (1997) and S. L. Brown et al. (2009) have shown the potential benefits of social closeness on self-reported measures, behavioral measures and hormonal correlates (progesterone) of prosociality. We could not replicate these results in a laboratory environment (A. Aron et al. (1997) used a more ecological environment: in a classroom) and importantly with the experimenter blind to the treatment condition (S. L. Brown et al. (2009) do not report that the experimental design was double blind). Moreover, a large part of the effect obtained by S. L. Brown et al. (2009) was due to a diminution of progesterone level in the control condition which was not similar to A. Aron et al. (1997) (while the social closeness condition was the same as A. Aron et al. (1997)). Overall, our results do not confirm the prosocial benefits nor the efficiency of social closeness manipulation.

Our data highlights the importance of experimental settings in the effects observed after interpersonal interactions. With this study, we show that it is possible to automatically program experimental manipulation of interpersonal relationships, which, to our knowledge, has never been done before. This paradigm was expected to test causal predictions concerning the influence of social interactions on physiological flexibility (Kok & Fredrickson, 2010; Porges, 2007). However, we can observe that results obtained without these methodological precautions (S. L. Brown et al., 2009) can not be reproduced here. As a consequence, this questions whether short-term positive social relationships actually impact physiological flexibility. Data from the current study does not support this causal pathway from the social to physiological levels of the polyvagal theory (Porges, 2007). As reported by Shahrestani et al. (2015), very few study attempted to test this causal relationship, and when reporting evidence toward a causal nature of the polyvagal theory, methodological biases (S. L. Brown et al., 2009; Kathi J Kemper & Shaltout, 2011; Willemen et al., 2008) can lead to question the conclusions. Here we show that the interplay between prosociality and heart-brain interactions proposed in the polyvagal theory (Porges, 2007) cannot be explained by the causal role of prosociality.

Several research directions can be explored in order to further explore this question. Our data do not show evidence toward a causal pathway in the polyvagal theory (Porges, 2007), but it worth to decline this kind of paradigm in order to test important modalities influencing social closeness.Indeed, the polyvagal theory suggest an association between sociability and autonomic flexibility allowed by efficient heart-brain interactions. In order to test the causal direction of this theory, it is needed to examine whether manipulating sociability does influence heart-brain interactions. Our study suggests that short term double-blind sham-controlled conditions run against this claim. We propose to test longer social closeness generation protocols in future experiments in order to test the importance of interaction time to influence physiological flexibility. Indeed, it is possible that the amount of retroactions between heart-brain functioning and inter-individual experiences has to be more frequent to be inserted in – even slightly – maintained interactions (Balliet & Van Lange, 2013; Boyer, Firat, & Leeuwen, 2015; Keltner, Kogan, Piff, & Saturn, 2014; Porges, 2007; S. C. Walker & McGlone, 2013). As a consequence, longer times or several repetitions of social interactions might be necessary to observe a physiological effect at the level of HF-HRV. If carried-out in rigorous methodological conditions, such a protocol could give further insight about the nature of prosocial processes involved in the polyvagal theory (Porges, 2007).

The proposition of Kok & Fredrickson (2010) concerning the bidirectional interplay between autonomic flexibility and prosociality can corroborate the idea that long-term laboratory manipulation might help to test the polyvagal theory (Porges, 2007). Indeed, the learning mechanisms involved in the management of unsafety and uncertainty in the environment (Brosschot et al., 2016b, 2016a) are likely to be inherently dependent on a minimum of interaction time with strangers in the social domain. For now, the polyvagal theory is not supported by our data in the social to physiological direction. Because we show the feasibility of rigorous methodological set-up, we suggest that further researchers aiming at testing the theory should carry-out experiments based on double-blind protocols.

### Conclusions

We aimed to test a possible causal pathway of the polyvagal theory (Porges, 2007) according to which prosocial behaviors can positively impact heart-brain dynamics. Our data does not support the theory in this direction. We suggest that further studies attempting to test the theory should focus on rigorous methodological features such as double-blind design protocols.

## Aknowledgements

We thank Stefan Agrigoroaei for technical support in data collection. We also thank Anthony Lane, Anne Kever, Julie Terache, Bertrand Beffara, Anne Weisgerber, Anael Le Runigo, and Emmanuel Daveau for their work on the translation of the social closeness generation procedure. We thank fabrice Damon and Ladislas Nalborczyk for their useful comments and constructive remarks. This research was funded by the French CNRS and the Belgian FNRS.

## Footnotes

1 No transformation allowed to get normality distribution on reaction times

